# Flow-wise or path-wise: diffusion in a fragmented dendritic network and implications for eels

**DOI:** 10.1101/323006

**Authors:** J. Domange, P. Lambert, L. Beaulaton, H. Drouineau

## Abstract

The European eel (*Anguilla anguilla*) is a catadromous species that reproduces at sea and inhabits continental waters during its growth phase. River fragmentation due to obstacles is considered one cause of the decline of this species. However, the colonization process of river catchments by eels is still poorly understood. In this article, we compare two scenarios for the diffusion of eels within river catchments: a path-wise scenario, in which movements are totally random, and a flow-wise scenario, in which movements are partially oriented. Based on these two scenarios, we attempted to predict the distribution of eels within dendritic river catchments, explicitly accounting for the presence of obstacles to movement. The model was fitted to a long-time series of electro-fishing data. The results suggest that the path-wise scenario is more predominant than the flow-wise scenario. Moreover, results show that the distribution of eels in river catchment depends on (i) the types of movements carried out by eels, (ii) the configuration of river networks and (iii) the positions of obstacles within catchments.

## 1 Introduction

Movement is a key feature of living organisms (Nathan et al., 2008), allowing them to avoid predation, feed, or mate or, more generally, to find suitable habitats and to undergo key phases of their life cycle. Dingle and Drake (2007) distinguished different types of movements. A first type of movement takes place within an individual’s home range, often referred to as “station-keeping” (Dingle, 1996), which generally corresponds to foraging, predation avoidance or defense of territory and implies simple orientation mechanisms and short time scales. The second type of movement involves an individual changing its home range and can be divided into ranging or migration (Dingle and Drake, 2007). Ranging corresponds to the search for specific resources (e.g., food, social interactions) and generally stops when the resource is obtained (Jeltsch et al., 2013). On the other hand, migration is generally a response to environmental cues, such as temperature or photoperiod, rather than fluctuations in resources and the availability of mates (Dingle, 2006). This type of movement applies to most individuals in the population and implies the existence of a return journey (Dingle, 2006; Dingle and Drake, 2007). Additionally, it occurs on longer time scales, requires elaborate orientation mechanisms, and is particularly prevalent in the animal kingdom (Wilcove and Wikelski, 2008). Interestingly, most migratory animals, whatever their taxon, are suffering dramatic declines because of global changes, especially over-exploitation, climate change and ecosystem fragmentation (Berger et al., 2008; McDowall, 1999; Sanderson et al., 2006; Wilcove and Wikelski, 2008).

The disruption of ecological connectivity due to ecological fragmentation is currently one of the major challenges in ecosystem restoration and conservation (Crook et al., 2009; Drouineau et al., 2018; Humphries and Winemiller, 2009; Kondolf et al., 2006; Sutherland et al., 2013; Tischendorf and Fahrig, 2000a, 2000b). Because dams and weirs have modified flow regimes and, thus, habitats, resulting in the impairment of fish movements and gene flow, human-induced river fragmentation has had visible impacts on fishes at the scale of assemblages (Branco et al., 2012; Santos et al., 2013), species/metapopulations (Horreo et al., 2011; Wollebaek et al., 2011) or populations (Limburg and Waldman, 2009). Among fishes, diadromous fishes divide their life-cycle between marine and continental habitats (McDowall, 1968; Myers, 1949). Similar to other migratory species in other taxa, fishes have suffered important declines throughout the world (Limburg and Waldman, 2009; McDowall, 1999), and many of them are now considered vulnerable or in danger of extinction. Though many anthropogenic pressures are involved in these declines, dams are noted as a key factor (Limburg and Waldman, 2009) responsible for the extirpation of various populations in river basins, or their confinement in restricted areas of river basins (Coutant and Whitney, 2000; Fukushima et al., 2007; Kondolf, 1997; Larinier, 2001; Limburg and Waldman, 2009; Porcher and Travade, 1992).

In temperate zones, we generally distinguish two types of diadromous species (McDowall, 1988). Anadromous species, such as many salmon species, reproduce in freshwater and grow at sea. Consequently, river fragmentation due to anthropogenic barriers impairs their downstream migration from their spawning grounds to their growth habitat and their return migration to the spawning grounds, thus impairing the completion of their life cycle. On the other hand, catadromous species, such as all temperate eels (*Anguilla anguilla*, *A. rostrata* and *A. japonica*), reproduce at sea and grow in continental waters. Thus, for these species, human barriers can also impair both upstream (Drouineau et al., 2015; Mouton et al., 2014; Piper et al., 2012) and downstream migration (Acou et al., 2008; Drouineau et al., 2017; Winter et al., 2007), in addition to limiting their movements during the growth phase, such as ranging or station-keeping. The situation is even more complex for temperate eels because they exhibit wide plasticity in their use of continental habitats (Daverat et al., 2006; Tsukamoto et al., 1998) and are able to settle in brackish estuaries or upstream habitats. Therefore, it is difficult to assess the impact of barriers on such species because it is difficult to determine how many barriers they will have to pass through. Moreover, the dendritic configuration of river catchments may lead to complex interactions between obstacles (Kuby et al., 2005; O’Hanley and Tomberlin, 2005; Palmer and Bernhardt, 2006) though some mathematical simulations tend to show the opposite (Padgham and Webb, 2010; Webb and Padgham, 2013). Consequently, quantifying the impact of obstacles on catadromous species requires a better understanding of the movement process in a dendritic river network.

The three species in this group reproduce at sea (Béguer-Pon et al., 2015; Schmidt, 1923; Tesch, 2003; Tsukamoto, 1992), after which their larvae, referred to as leptocephalea, are transported to continental shelves via a long passive drift (Bonhommeau et al., 2009), where they become glass eels. The eels then enter continental waters, where they become pigmented yellow eels, and remain there during their growth phase, which lasts several years. After a period of between 3 and 30 years, yellow eels metamorphose into silver eels, migrate back to the sea, and travel across the ocean to their oceanic spawning grounds (Tesch, 2003). These three species have suffered a dramatic decline since the late 1970s/early 1980s (Dekker, 2009; Dekker and Casselman, 2014), and they are now classified as endangered (Jacoby et al., 2014; Jacoby and Gollock, 2014a) or critically endangered (Jacoby and Gollock, 2014b) by the IUCN. In this context, the European Union has enforced an eel-specific Council Regulation (N°1100/2007) requiring a reduction of anthropogenic mortality, including dams, along with measures to facilitate upstream and downstream migration. Moreover, this Regulation sets a management target for Member States: the escapement of silver eels from inland waters should be superior to 40 % of the escapement in conditions without any anthropogenic influences. Consequently, tools are needed to estimate pristine escapement and to assess the impact of barriers on current escapement, and such tools require a better understanding of eel movements in continental waters.

Regarding the physiological mechanisms implicated in glass eel upstream migration, Imbert et al. (2010) demonstrated the role of thyroidal hormones in the shift between migratory and resident behavior. Bureau du Colombier et al. (2008) suggested that the migratory behavior of these eels could also be related to their feeding behavior, as eels with a high propensity to migrate also exhibited rapid resumption of feeding upon entering continental waters. From an evolutionary perspective, Edeline (2007) proposed that habitat selection would be the result of a conditional evolutionarily stable strategy (Gardner et al., 1987) in which habitat selection by individuals would be a trade-off between mortality reduction in upstream habitats (though this was also discussed by Cairns et al. (2009)), higher growth rates in downstream habitats and energy costs linked to migration. Drouineau et al. (2014) demonstrated that patterns of eel distribution within river catchments could result from fitness optimization, especially when taking into account intra-specific competition. Part of this pattern may also have a genetic basis with spatially variable selection (Gagnaire et al., 2012; Ulrik et al., 2014). Based on analyzing yellow eel density estimates in different river catchments, Ibboston et al. (2002) demonstrated that the density of eels from downstream to upstream habitats follows a Gaussian distribution. He concluded that whatever the underlying mechanism, the emergent movement process can be modeled as a diffusion process, which is a type of process that is widely used to model movements in ecology (Okubo and Levin, 2013). Naively, a diffusive movement corresponds to a random movement that is not orientated by any gradient. Diffusion of individuals leads to their progressive dispersion and to a Gaussian spatial distribution. However, two problems arise in association with diffusion. First, the dendritic nature of river catchments raises the question of the behavior of fishes at river junctions. More specifically, Johnson et al. (1995) showed that different options for modeling fish behavior at river junctions, which they referred to as path-wise diffusion or flow-wise diffusion, based on different ecological assumptions, lead to significantly different distributions of fishes in a dendritic river network.

Under path-wise diffusion, individuals explore all available habitats; at a river junction, they will select any available reach with similar probability. This scenario would correspond to an exploration directed toward resource seeking without any effect of river flow. On the other hand, under an unbiased flow-wise diffusion, at a river junction half of individuals move downstream and half upstream, after which they select a reach among the available reaches (flow-wise partitioning means that the probabilities of moving upstream or downstream remain constant whatever the configuration). This scenario results in an oriented exploration of the catchment. Second, eels do not move in a free river network but in a fragmented river network, i.e. in a river network in which movements are partially impeded by dams or weirs, which modifies the diffusion process. Moreover, the impact of barriers is likely to be distinct between flow-wise and path-wise diffusion scenarios.

In this context, we developed a model referred to as TABASCO (spaTialised Anguilla BASin COlonization assessment model) to explore the consequences of flow-wise versus path-wise diffusion for eel colonization, along with the subsequent impact of obstacles under the two scenarios. Diffusion is represented using either a path-or flow-wise random walk between successive river reaches. The model was applied to the French hydrographic networks, which are disrupted by human obstacles, and fitted to electrofishing data from 1990 to 2009.

## 2 Materials and methods

### 2.1 Data

Several georeferenced databases were used in this study: RHT (http://www.irstea.fr/en/rht) (Pella et al., 2012), BDMAP (Base de Donnée sur les Milieux Aquatiques et Piscicoles) (Poulet et al., 2011) and the “Référentiel des Obstacles á l’Ecoulement” (ROE ^®^ database, finalized version 6.0, http://www.onema.fr/REFERENTIEL-DES-OBSTACLES-A-L).

RHT (Fig. 1) is a theoretical representation of the French hydrological network, composed of 114,600 reaches. Several important hydrological features are available for each reach, such as the average discharge, width, height, and flow duration curves.

**Fig. 1.**
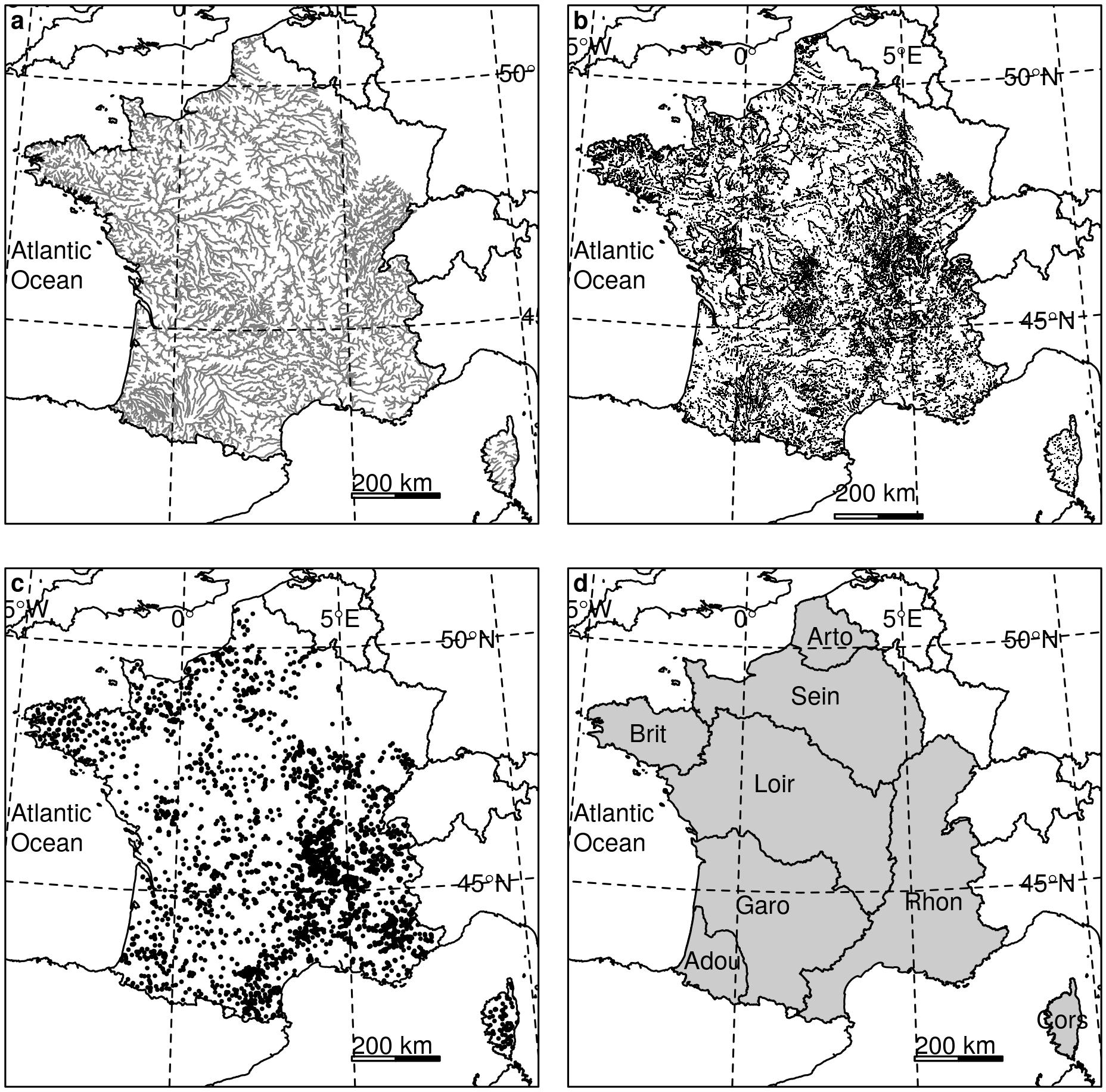
Hydrographic network used in the model (a; only reaches that are more than 5 m wide are plotted=, adapted from RHT (Pella et al., 2012). Maps of the obstacles (b), fishing operations (c) and Eel Management Units (d; Brit=Brittany, Arto=Artois Picardie, Sein=Seine, Loir=Loire, Adou=Adour, Garo=Garonne-Dordogne-Charente-Seudre, Cors=Corsica, Rhon=Rhône-Méditerranée) used in this study.

ROE is an inventory of all obstacles to river flow in France. In the 6^th^ version, approximately 70,000 obstacles were included in the database. We restricted the dataset to transverse obstacles of weir and dam types and removed auxiliary obstacles. The remaining obstacles (~60,000) were projected onto the RHT reaches (Fig. 1b), which were subsequently separated at the level of obstacles. Reaches longer than 2 km were then split again, so that the final reaches were all shorter than 2 km.

The resulting hydrographic network (Fig. 1a) was composed of ~225,000 reaches with corresponding hydrological attributes.

BDMAP stores the results of electro-fishing operations in the French territory. We selected all complete two-pass electro-fishing operations from 1990 to 2009 and projected them onto our hydrographic network (Fig. 1c – 8531 operations), so that we were able to attribute a reach to each fishing operation. The Carle and Strub method (Carle and Strub, 1978) was used to estimate the number of eels at the sampling station and the associated standard error for each electrofishing operation. We used abundance estimates for all yellow eels with total lengths of 150 mm to 900 mm, i.e. yellow eels, though smaller eels may display different behaviours than larger and older eels. Indeed, an ontogenic shift from migratory to sedentary behaviours occurs at about 200 mm (Imbert et al., 2010). Data are presented in supplementary material.

Finally, only catchments (and corresponding fishing operations and obstacles) located entirely within France were retained in our analysis because the entire hydrographic network was necessary to carry out the random walk from the outlet (in this study, outlets corresponds to estuaries, i.e. where rivers meet the sea). Consequently, many catchments in the northern and eastern parts of France (which flow into Belgium and the Netherlands) were removed, and only 8 Eel Management Units (EMUs) were considered (Fig. 1d). The catchments in each EMU are assumed to exhibit a certain level of homogeneity regarding eel ecology and anthropogenic pressures.

Figure 2 illustrates the contrast in both hydrographic network configuration and position of obstacles between two EMUs. Hydrographic networks are more ramified in Adour than in Artois-Picardie that display more linear networks. Moreover, a large number of obstacles are located close to the sea in Artois-Picardie whereas most obstacles are located in upstream reaches in Adour.

**Fig. 2.**
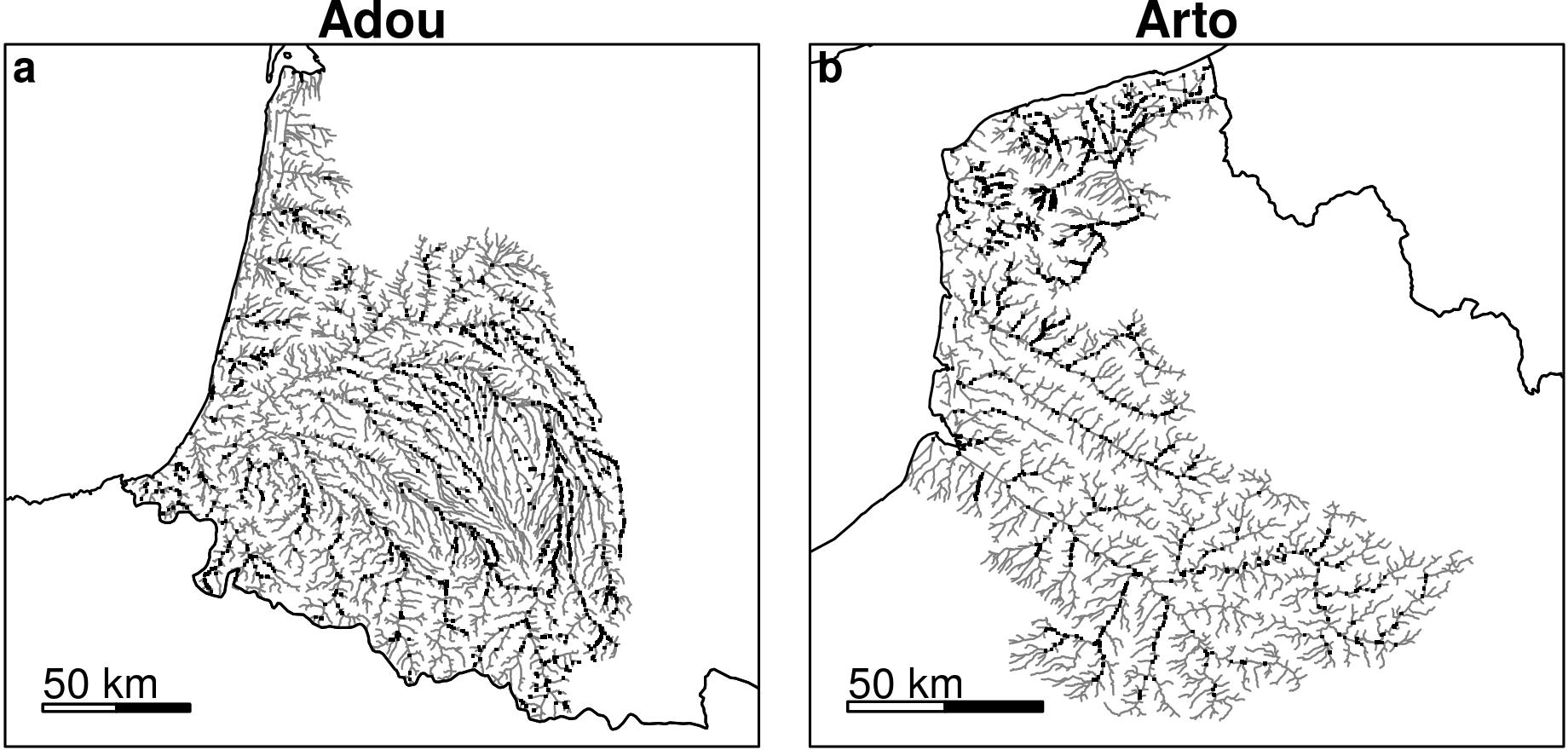
Hydrographic networks (grey lines) and obstacles (black points) in two EMUs (a Adour, b Artois-Picardie).

All geographic operations were carried out using PostGIS, which is a software that adds support for geographic objects to PostgreSQL, an object-relational database management system. More specifically, the river network was represented and divided using a topology object type.

### 2.2 Model

#### 2.2.1 Notations

A summary of the model parameters and the notations used in this study is presented in Table 1.

**Table 1.**
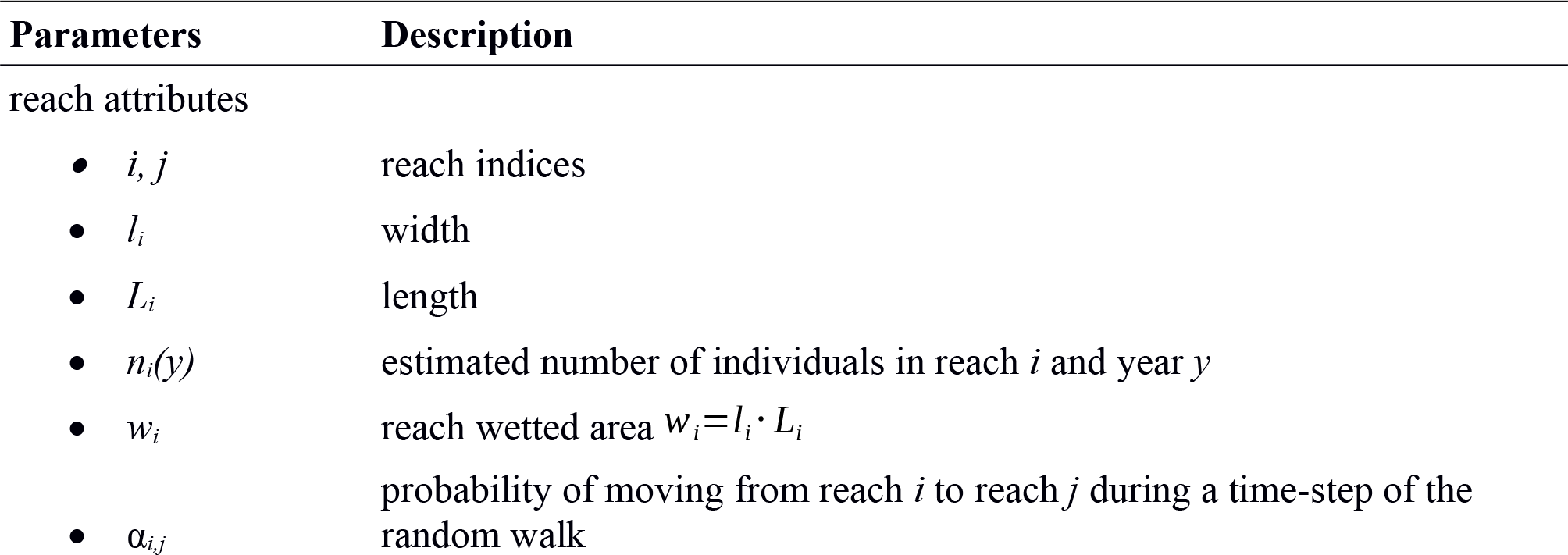
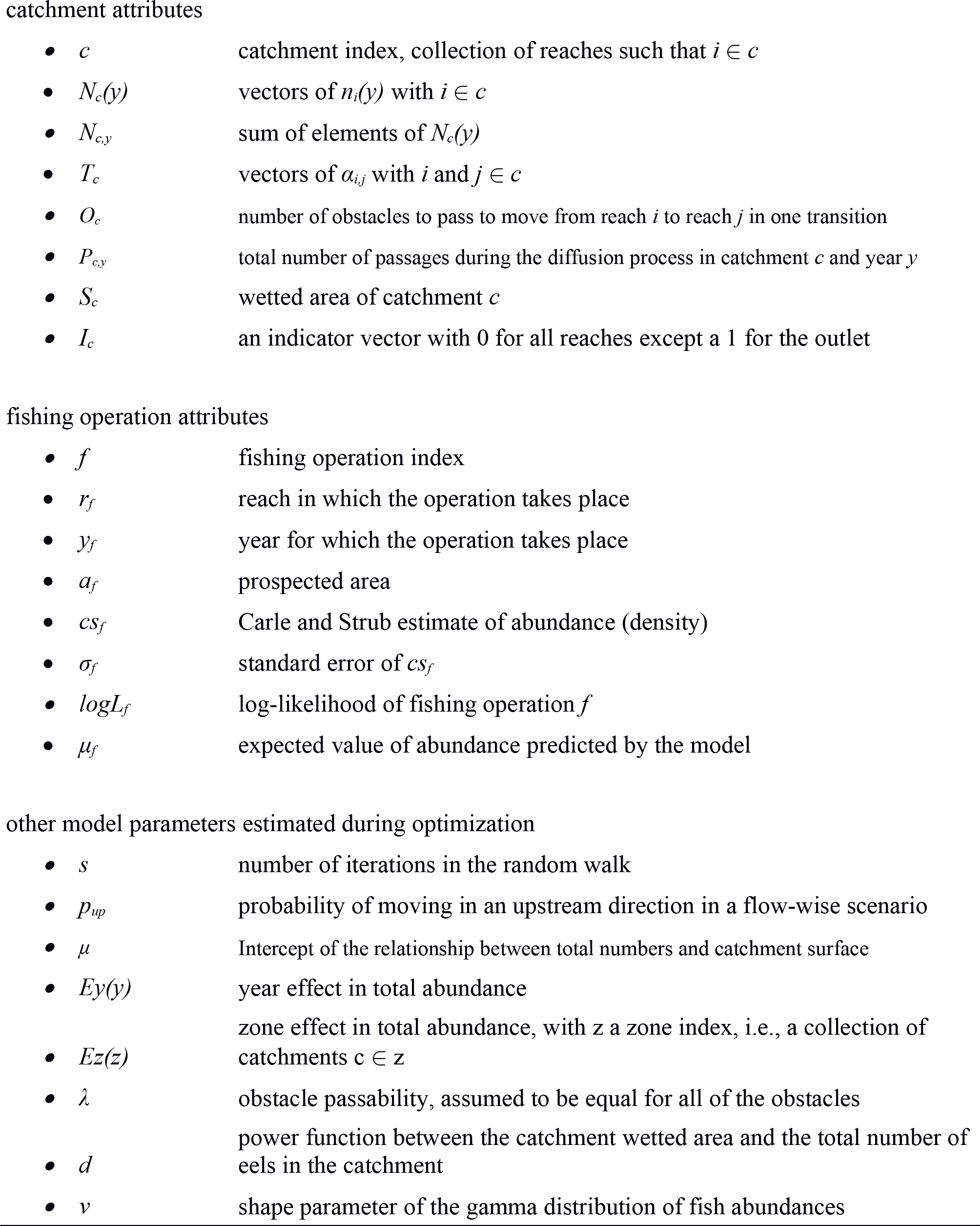
Problem-solving strategies that separate qualitative/pictorial steps from mathematical steps.

#### State model: transition matrix

Eel movement was modeled via a random walk (DeAngelis and Yeh, 1984; Pearson, 1905). This random walk was implemented through the construction of a transition matrix for each river catchment that included the probabilities of moving from a reach to its adjacent reaches during an iteration step. In a pure isotropic and unidimensional random walk, the probability of moving in the reach upstream would be 0.5, and the probability of moving in the reach downstream would be 0.5. However, in a catchment, reaches do not exhibit the same lengths or the same wetted area (which is used here as a proxy of available habitats) and are connected according to the river network topology. We also have to account for obstacles that hamper movements. To do so, we consider that at each iteration, an eel in reach *i* in catchment *c* can reach all *j* reaches from *c* if *i* and *j* are adjacent or if the distance between *i* and *j* is less than 1 kilometer. This limit of 1 kilometer was set to avoid anomalies due to very short reaches (67% of reaches were between 1 and 2 kilometers long). The computation of transition probabilities is summarized in Fig. 3. Briefly, the probability of moving between reaches depends on their respective wetted areas and the passability of existing obstacles. In a flow-wise scenario, individuals first choose a direction (with probabilities *p*_*up*_ and 1-*p*_*up*_), after which the reach probability of moving to a reach depends on its wetted area with respect to wetted areas of reaches in that direction. In a path-wise scenario, individuals can move in any reach independently of the direction. Individuals that are blocked (either because of an obstacle or because there are no reaches available upstream or downstream) are assumed to remain in their reach.

**Fig. 3.**
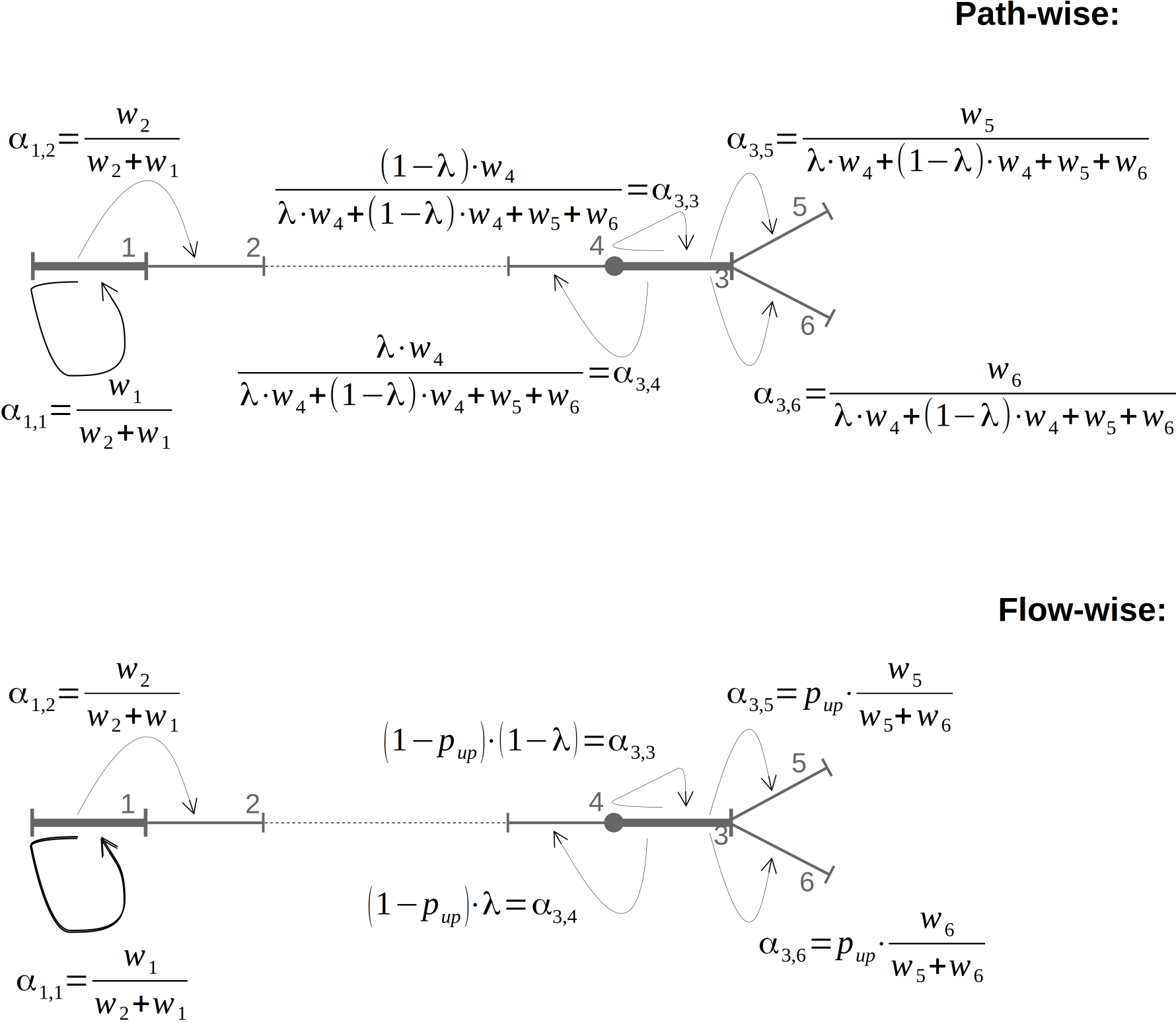
Computation of the probability of transition from two reaches (in bold) to adjacent reaches (thick segments) according to the scenario. A circle represents an obstacle. The river is assumed to flow from the right of the figure towards its left. Reach 1 is an outlet so there are no reaches downstream. Weights *w*_*i*_ are based on reach hydro-morphological characteristics and are known without errors. *λ* (passability) and *p*_*up*_ (proportion of individuals moving upstream) are estimated during the optimisation procedure.

**Fig. 4.**
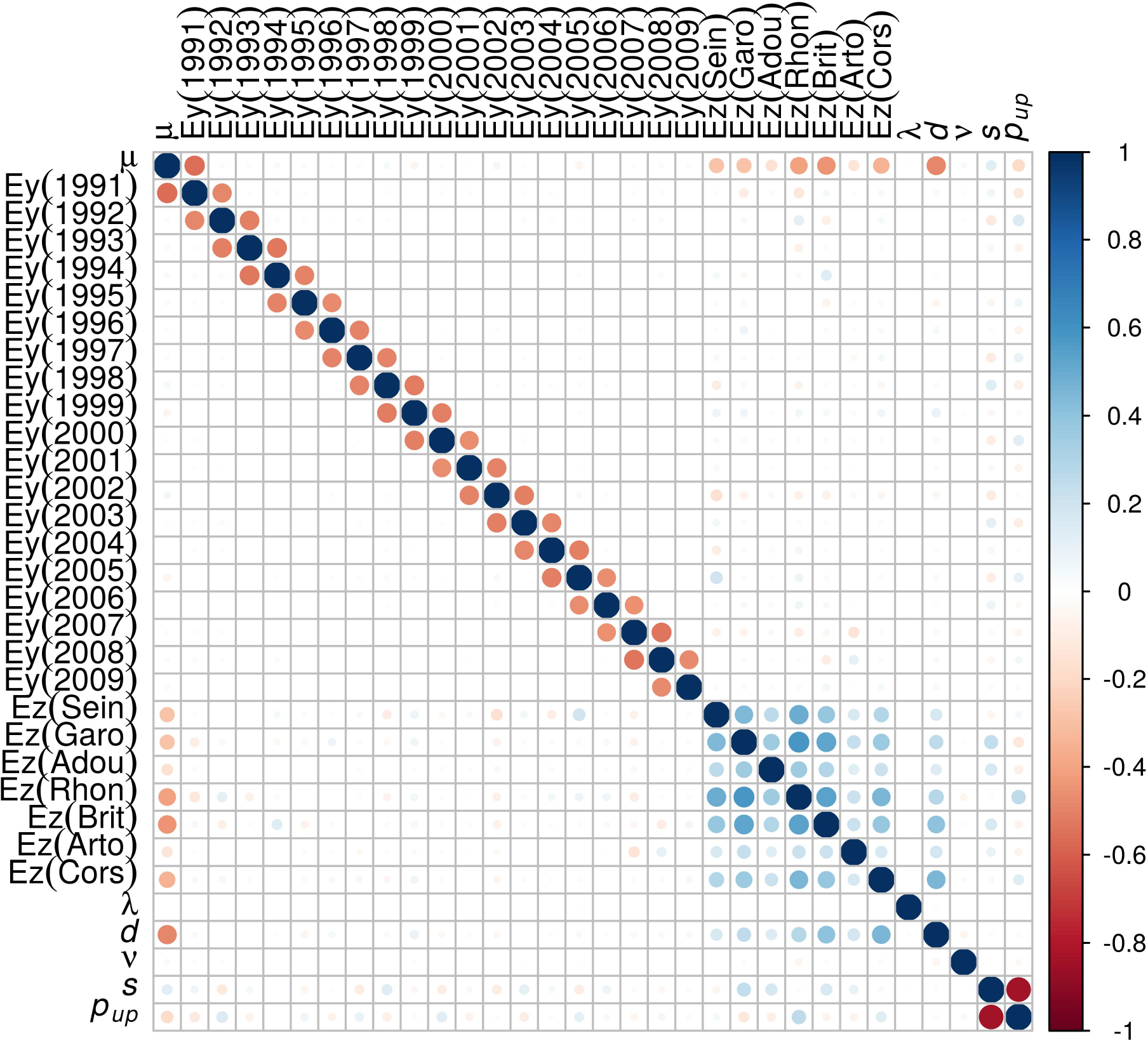
Correlation matrix of parameter estimates for the flow-wise scenario

**Fig. 5.**
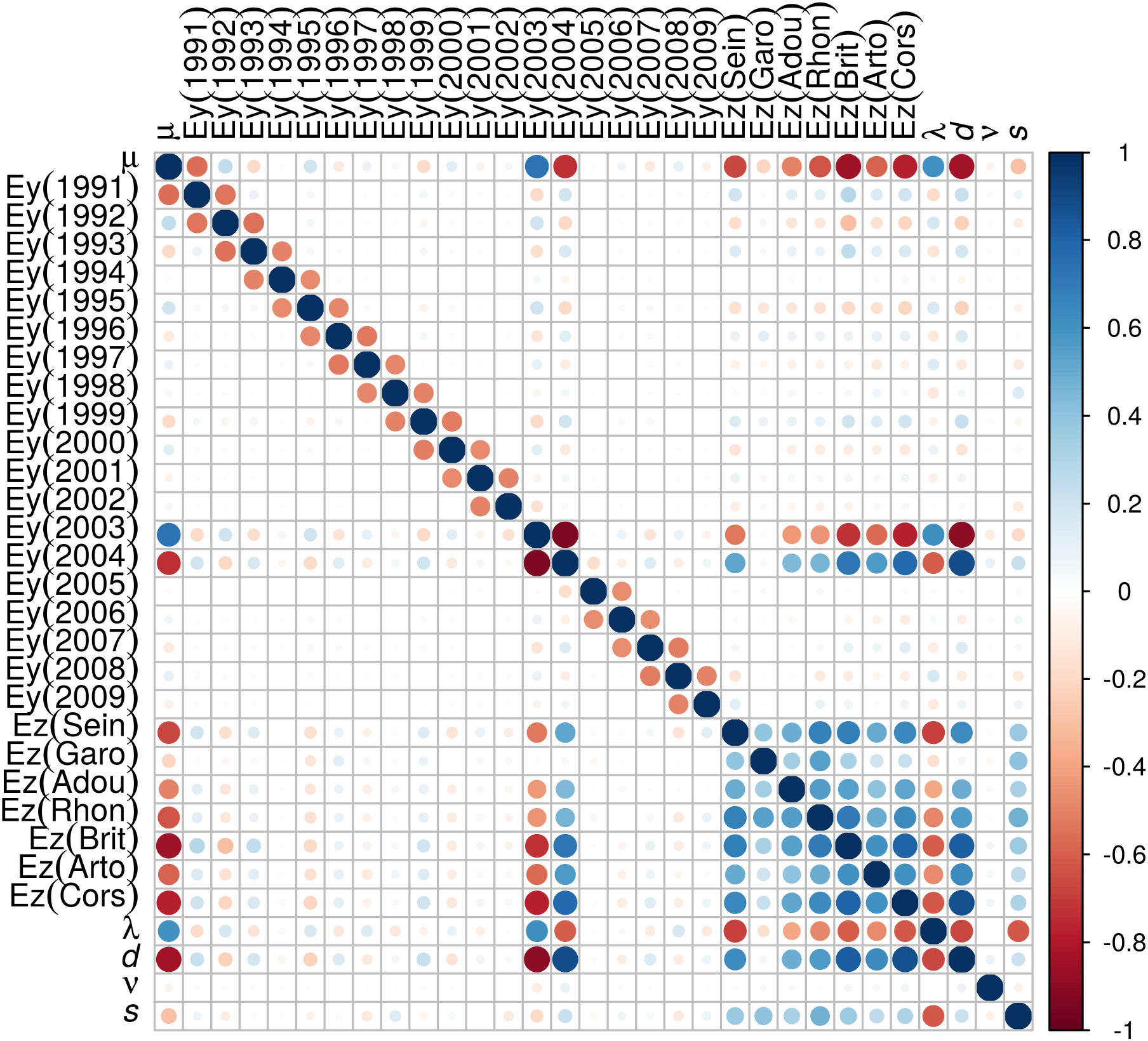
Correlation matrix of parameter estimates for the path-wise scenario.

The transition matrix, *T*_*c*_, stores the probability of moving from one reach (in columns) in catchment *c* to another (in rows).

The column vector of individual numbers [*n*_*i*_(*y*)]_*i*∈*c*_ in each reach (*i*) of catchment *c* for year *y* is calculated as follows:

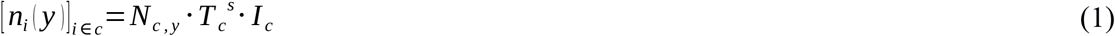

where *Ic* is a column vector of zeros, except for a value of one corresponding to the outlet reach.

Therefore, 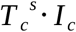 mimics a *s*-step random walk with individuals originating from the river outlets (i.e. from the sea) and provides the proportions of individuals in each reach of the catchment. *N*_*c,y*_ represents the total number of eels in catchment *c* for year *y*. *N*_*c,y*_ was considered to be the product of a year effect (*Ey(y)*), a zone (corresponding to an EMU) effect (*Ez(z)*) and a power function of the wetted areas of the rivers in the catchment (*S*_*c*_):

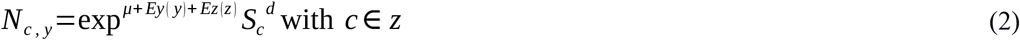

The year effect corresponding to 1990 and the EMU effect for the Loire EMU were set equal to zero to preserve model identifiability (i.e. they correspond to our baseline, as in classical regression with categorical predictors). We used a power function since catchment discharge increases with catchment wetted area as a power function (Burgers et al., 2014), and river discharge is thought to attract larvae (Crivelli et al., 2008; Drouineau et al., 2016; Elie and Rochard, 1994).

The total number of obstacles passages during the whole random walk is given by

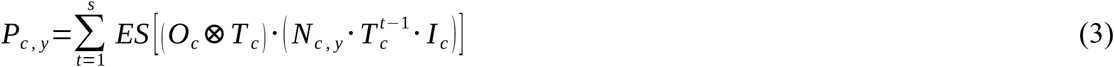

with ⊗ the matrix element-wise product and ES denoting the sum of all the elements of a matrix.

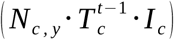 is the vector containing the number of individuals in each reach at step *t−1*, which multiplied by the proportion of individuals moving to a reach and by the number of passed obstacles during this transition, gives a matrix containing the number of passages for all transitions.

#### 2.2.3 Observation model

The model was fitted by comparing the densities estimated by electrofishing operations with the densities predicted by the model in corresponding reaches.

For a fishing operation (*f*) taking place in catchment *c* in reach *r*_*f*_ during year *y*, the Carle and Strub estimator of abundance (*cs*_*f*_) and the associated standard error (σ_*f*_) were available. Since the Carle and Strub method is based on a likelihood procedure, the 95 % confidence interval of *cs*_*f*_ was between cs_*f*_ −1.96 · σ_*f.*_. and cs_*f*_ + 1.96 σ_*f*_ (large samples approximation assuming that likelihood assumptions were met). If the fishing station has a surface *a*_*f*_, the model predicted that the expected number of eels was 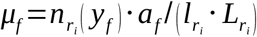, i.e. the number of fishes predicted by the model in the reach 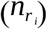 multiplied by the proportion of the reach that was sampled during the fishing operation. Because eels are not uniformly distributed within a reach, we assumed that the number of fishes caught during a fishing-operation follows a gamma distribution of mean *μ*_*f*_ with a constant shape parameter (*v*) that governs the level of spatial heterogeneity (a low v increases the heterogeneity). A gamma distribution is often used in fisheries sciences to model abundance indices with overdispersion (Drouineau et al., 2010; Froysa et al., 2002), which can occur because of aggregative behavior of fishes and spatial heterogeneity of habitats. The description of the gamma distribution can be found in McCullagh and Nelder (1989). The log-likelihood is then as follows:

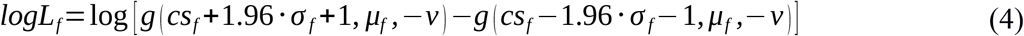

where *g* is the cumulative distribution function of the gamma distribution. We added +1 and −1 to the boundaries of the confidence intervals because the Carle and Strub method is based on integer optimization, so that the true value can be included between *CS*_*f*_−1 and *CS*_*f*_+1 and to avoid over-fitting of the data for which σ_*f*_ is too small (a null σ_*f*_ sometimes arises because of this integer optimization). The log-likelihood in equation 4 corresponds to the log-likelihood of a double-censored data. By doing this, we account for the uncertainty around the Carl and Strub estimator, quantified by its error. Rather than using the probability density function at a precise but uncertain point, we preferred to measure the probability that the estimate is within the range of the confidence interval. Using the cumulative distribution function instead of the density function is frequently used to fit survival models (Cox and Oakes, 1984; De Gruttola and Lagakos, 1989), and in many kind of environmental and ecological models (Frome and Wambach, 2005; Helsel, 2005; Munoz et al., 2017), by maximising the likelihood in the presence of poorly measured data, for example data above or below a detection threshold. By doing this, we consider that all values in the confidence interval are equally likely and therefore we loose a little bit of information. However it allows to use the same function for both positive and null values and therefore to avoid to have two components in the likelihood function (the gamma distribution is not defined for null values) and to look for appropriate weights.

#### 2.2.4 Model implementation

The model was coded in C++. River catchments were implemented using graphs from the Boost C++ libraries (http://www.boost.org/). Matrix computations were carried out using two different libraries: Eigen (http://eigen.tuxfamily.org/index.php?title=Main_Page) (especially the implementation of the sparse transition matrix) and many statistical functions from Autodif (Fournier, 1996). Communication between the PostreSQL/PostGIS database and the C++ code was carried out using the pqxx library (http://pqxx.org/).

#### 2.2.5 Model fining

The model was fitted by maximizing the log-likelihood. Thirty-one parameters were estimated in this procedure (Table 1): year effects (*Ey(y)*) (20 parameters from 1990 to 2009), zone effects (Ez(z)) (7 parameters; there are 8 French EMUs, but the Loire EMU effect was set to zero and was used as a reference), obstacle passability (*λ*), the shape of the gamma distribution (*v*), the power parameter (*d*) and the number of transitions (*s*). Theoretically, parameter *s* is an integer, but we carried out relaxation by treating it as a continuous parameter to achieve optimization. A 32^nd^ parameter was estimated in the flow-wise scenario: *p*_*up*_.

The model was fitted using a CMA-ES algorithm (Hansen et al., 2003), which is an evolutionary metaheuristic algorithm, with default algorithm parameters.

After convergence, we computed the Hessian matrix to check the model convergence. The covariance matrix of parameter estimates, which was estimated as the inverse of the Hessian matrix, was used to compute confidence intervals around parameter estimates.

#### 2.2.6 Analysis of the results

For each scenario, the model predicts the number and density of eels for each reach and each year. Since the model is complex, analysing outputs is not straightforward. Therefore, we chose to fit a metamodel on model outputs. A metamodel (aka surrogate model) is a simple, generally statistical model that mimics the studied complex model in order to explore its properties. Metamodelling is commonly used to carry out sensitivity analysis (Faivre et al., 2013; Kleijnen, 2015; Saltelli et al., 2000) or calibration of complex models (Jones et al., 1998). Here, we fitted boosted regression trees (BRT) (Elith et al., 2006; Friedman, 2002) on model outputs to explore the relative influence of factors on outputs. BRTs combine traditional decision trees and boosting techniques, by combining multiple very simple decision trees to improve the overall predictive accuracy (Elith et al., 2008). In a regression context, BRT, similarly to traditional linear models, aims at predicting the value of an independent continuous variable based on continuous or categorical predictors. Because of their abilities to capture non-linearities and to account for complex interactions between factors (Elith et al., 2008), BRTs are very flexible and have been used to develop species distribution models in ecology (Elith et al., 2006; Lassalle et al., 2010). This flexibility makes them very relevant to be used as surrogate models of complex simulation models (Friedman, 2001). Interestingly, BRTs provide a measure of the relative importance of the predictors which quantifies the influence on each predictor on the considered output (Friedman, 2001).

We fitted two BRTs to address two specific questions. The first focuses on the influence of factors on the density of eels in a given reach. In this BRTs, we used the log density predicted by flow-wise scenario and path-wise scenario in each reach (for year 1999) as the output variable, and used characteristics of the reach, its position within the river catchment, characteristic of the catchment, its network configuration, the presence and densities of obstacles in the catchment and downstream the reach, and the type of movement (flow-wise of path-wise) as explanatory variables (see Table 2).

**Table 2.**
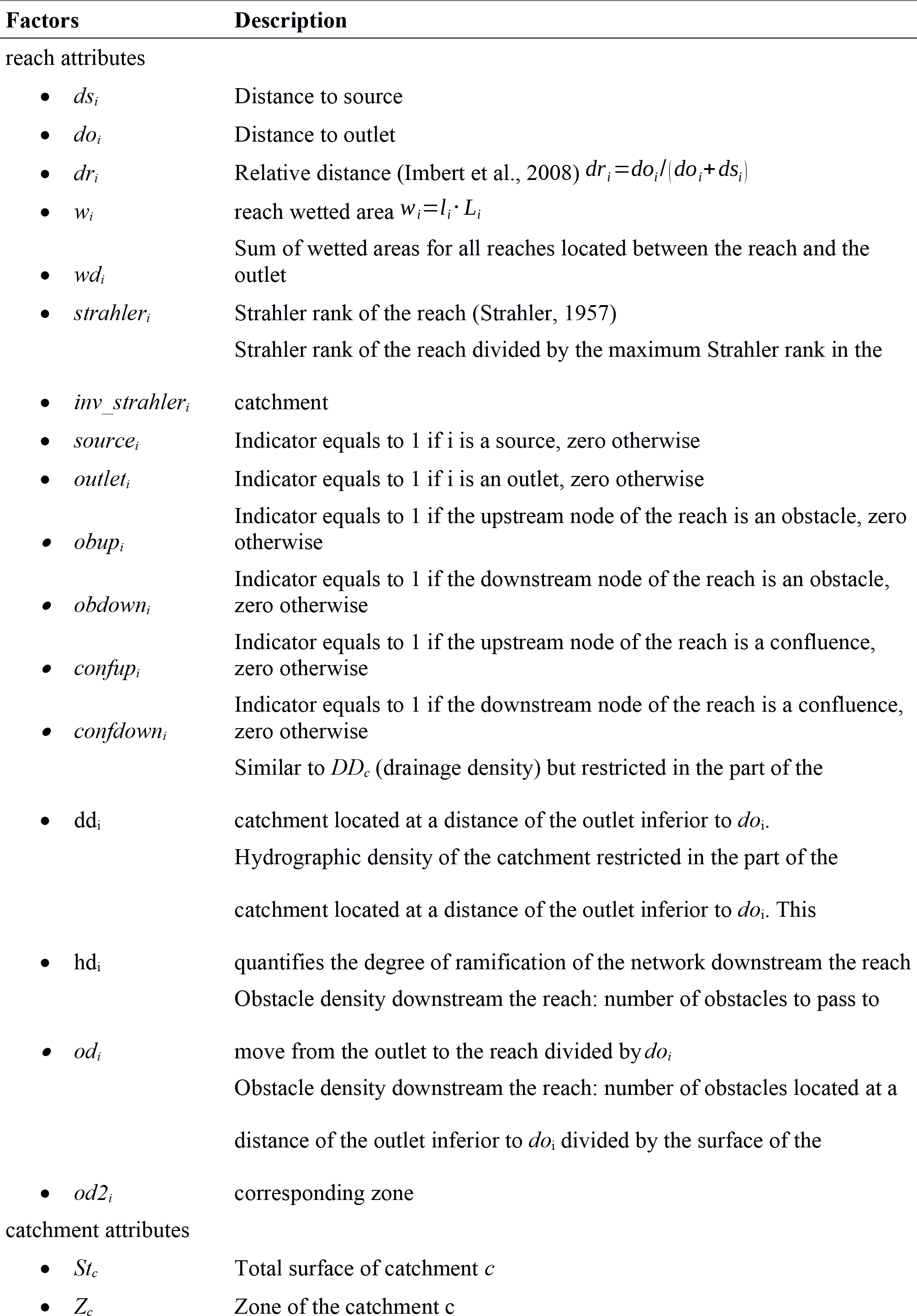
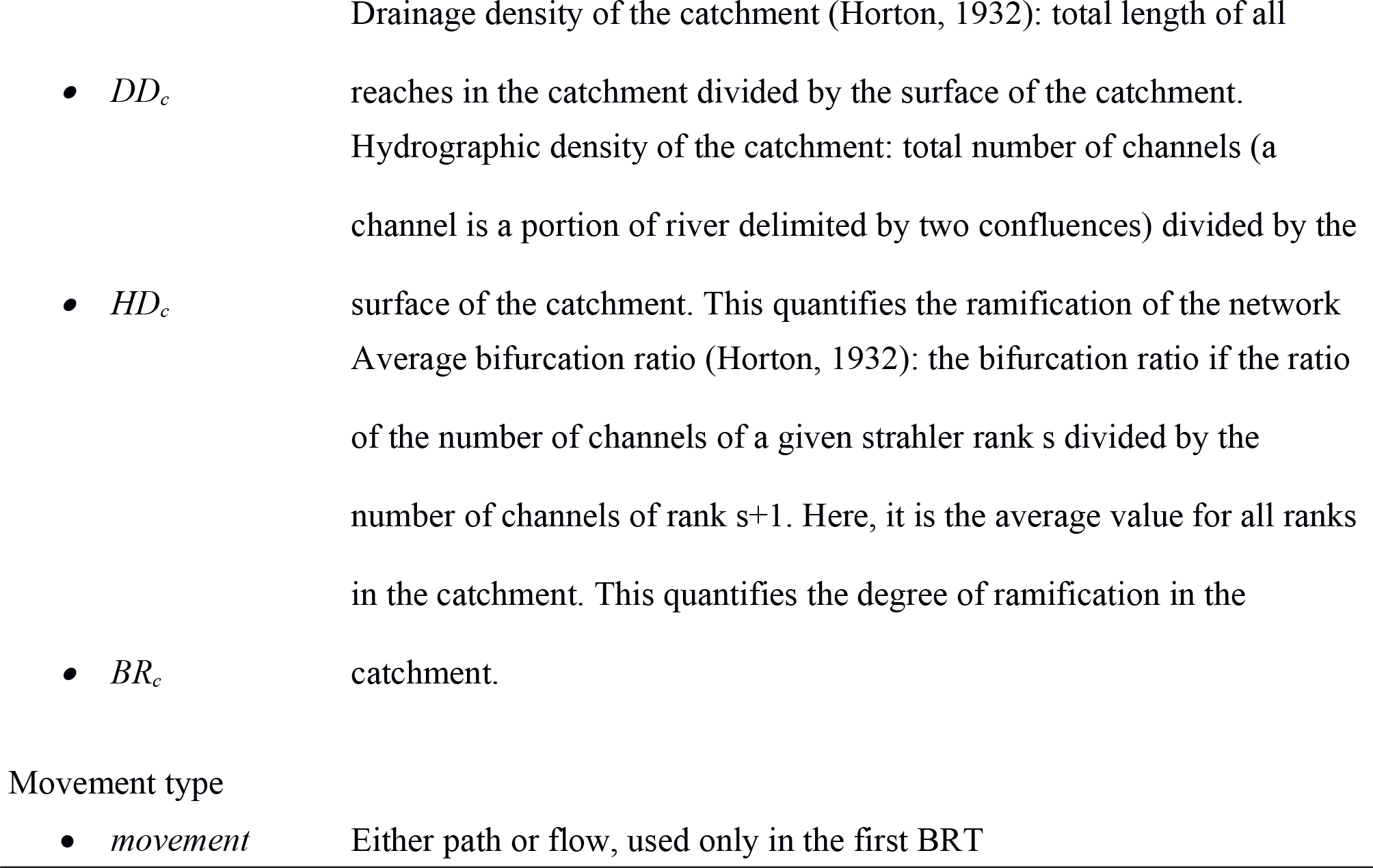
List of predictors used in the two BRTs.

The second BRT focused on the differences between flow-wise and path-wise scenarios. Here we used the difference of log densities in each reach (for year 1999) as the output variable and used the same explanatory variables as in the previous BRT, except movement type.

A Gaussian family was used for both BRTs and their main parameters (learning rates: 0.05, number of trees: 15.000 and tree complexity: 10) were set using a 10-fold cross-validation procedure, and we used a bagging fraction of 0.5, as recommended by Elith *et al.* (2008). BRTs were fitted using the package lightGBM in python (Ke et al., 2017).

## 3 Results

### 3.1 Model performance and estimated parameters

The model converged with both options (after 850 CMA-ES iterations and 11,900 model runs for the path-wise scenario; after 3,045 CMA-ES iterations and 42,630 model runs for the flow-wise scenario). Hessian matrices were positive definite for both scenario, confirming that the algorithm had indeed converged to a, at least local, optimum. However, strong correlations were observed among estimated parameters, especially for the path-wise scenario and EMUs effects.

The path-wise scenario led to a better log-likelihood than the flow-wise scenario (−10,095.1 versus −10,418.6). Given the numbers of parameters (31 and 32, respectively), the corresponding Akaike information criterion values were 20,252.2 and 20,901.1 for the path-wise and flow-wise options, respectively. This difference indicated that the path-wise scenario yielded a better fit to the data. Interestingly, the two scenarios produced different parameter estimates (Table 3). For example, the numbers of steps in the random walk process were different, with more steps in the path-wise scenario. Even the year effects and the EMU effects (*Ez(z)*) were different in the two scenarios. For example, the Adour and Artois-Picardie EMUs effects were significantly inferior to zero with the path wise scenario and not for the flow-wise scenario. For the year effects, years 1991, 1993, 2002 2005 and 2006 were significantly different from zero with a flow-wise scenario but not with a path-wise scenario. Conversely 1995 and 1997 had significant effects in the path-wise scenario but not with the flow-wise scenario. The difference in the *d* parameter indicated that the differences in abundance between catchments of different sizes were smoother in the flow-wise scenario compared with the path-wise scenario.

**Table 3.** Estimated values for model parameters and 95% confidence intervals in parenthesis. *Ez(Loire) and Ey(1990) were not estimated but fixed in the model.

| Parameter | Flow-wise | Path-wise |
| --- | --- | --- |
| $\mu$ | 14.29 (13.85,14.73) | 12.88 (12.19,13.57) |
| Ey(1990)* | 0 | 0 |
| Ey(1991) | 0.98 (0.49,1.47) | 0.42 (-0.04,0.88) |
| Ey(1992) | -1.64 (-2.13,-1.16) | -0.59 (-1.05,-0.14) |
| Ey(1993) | 0.6 (0.11,1.09) | -0.01 (-0.47,0.45) |
| Ey(1994) | -0.31 (-0.78,0.17) | -0.34 (-0.77,0.1) |
| Ey(1995) | 0.32 (-0.09,0.73) | 1.04 (0.64,1.44) |
| Ey(1996) | -0.44 (-0.82,-0.07) | -1.07 (-1.44,-0.71) |
| Ey(1997) | 0.33 (-0.04,0.69) | 0.61 (0.26,0.96) |
| Ey(1998) | -0.24 (-0.61,0.12) | -0.33 (-0.68,0.01) |
| Ey(1999) | -0.55 (-0.91,-0.18) | -0.31 (-0.66,0.04) |
| Ey(2000) | 1.07 (0.72,1.41) | 0.79 (0.47,1.12) |
| Ey(2001) | -0.07 (-0.4,0.26) | 0.14 (-0.18,0.45) |
| Ey(2002) | 0.48 (0.13,0.82) | 0.11 (-0.22,0.44) |
| Ey(2003) | -0.06 (-0.39,0.28) | -0.02 (-0.91,0.87) |
| Ey(2004) | -0.12 (-0.45,0.21) | -0.32 (-1.18,0.54) |
| Ey(2005) | -0.89 (-1.23,-0.56) | -0.28 (-0.6,0.03) |
| Ey(2006) | -0.45 (-0.78,-0.12) | -0.12 (-0.43,0.19) |
| Ey(2007) | 0.13 (-0.25,0.51) | -0.24 (-0.59,0.1) |
| Ey(2008) | 0.50 (0.07,0.92) | 0.78 (0.39,1.16) |
| Ey(2009) | -0.58 (-1.05,-0.12) | -0.71 (-1.13,-0.29) |
| Ez(Loire)* | 0 | 0 |
| Ez(Seine) | -0.24 (-0.47,-0.01) | -0.11 (-0.41,0.19) |
| Ez(Garonne) | -0.69 (-0.91,-0.47) | -0.84 (-1.05,-0.63) |
| Ez(Adour) | -0.28 (-0.66,0.11) | -0.84 (-1.25,-0.44) |
| Ez(Rhône) | -3.1 (-3.32,-2.88) | -2.45 (-2.69,-2.21) |
| Ez(Brittany) | -0.93 (-1.22,-0.65) | -0.87 (-1.33,-0.42) |
| Ez(Artois-Picardie) | -1.23 (-1.94,-0.51) | -0.89 (-1.69,-0.08) |
| Ez(Corsica) | -2.89 (-3.34,-2.44) | -1.68 (-2.43,-0.94) |
| $\lambda$ | 1 (1,1) | 0.5 (0.45,0.54) |
| $d$ | 0.42 (0.38,0.45) | 0.78 (0.67,0.89) |
| $v$ | 0.16 (0.17,0.15) | 0.19 (0.2,0.18) |
| $s$ | 18768 (17629,19980) | 27976 (26591,29433) |
| $p_{up}$ | 0.51 (0.51,0.51) | |

### 3.2 Distribution of eels in the territory-influence of factors on eel densities

The two scenarios produced contrasted results in terms of distribution in the territory (Fig. 6). With a path-scenario, eels tend to be more present in the main channel whereas they are more evenly listributed with a flow-wise scenario.

**Fig. 6.**
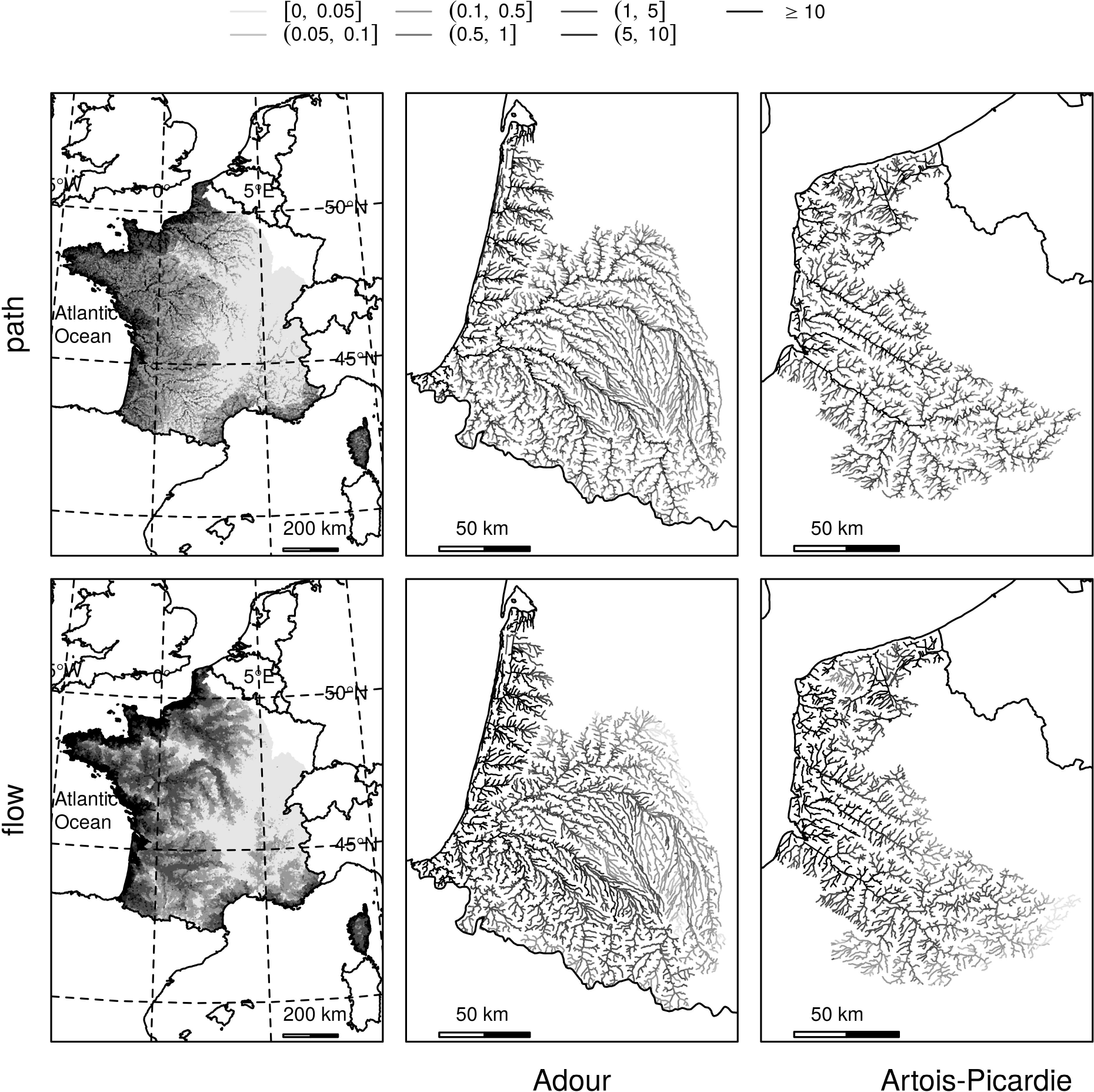
Density per reach (eels.100 m^−2^) under the path-wise (first line) and flow-wise (second line) scenarios for the year 1999, in France (first column), Adour (second column) and Artois-Picardie (third column). Maps for other years are presented in supplementary materials.

The analysis of the first BRT provides insights on the important factors that affect eel distribution. Eel density is mainly sensitive to reach attributes (Fig. 7). The distance to the sea (either *do*_*i*_ or *wd*_*j*_ which are difficult to disentangle because correlated) are unsurprisingly two very important factors. However, the number of obstacles to pass (*od*_*j*_ and also *od2*_*i*_) is also very important factor (the most important according to the BRT, but *do*_*i*_ or *wd*_*j*_ are correlated and both account for the distance to the sea). Other reach attributes influence the density in a reach, related to the quantity of available habitats (*w*_*i*_), the position in the network (*ds*_*i*_, *dr*_*i*_,) and the network configuration downstream the reach (hdi). Since the distribution of eels in the territory directly influence the number of obstacles that eel will have to pass during their downstream migration to the sea, this means that the overall impact of obstacles over silver eel depends on the type of movement, on the network configuration and the position of obstacles within the catchment. Attributes related to the river catchment have a minor influence.

**Fig. 7.**
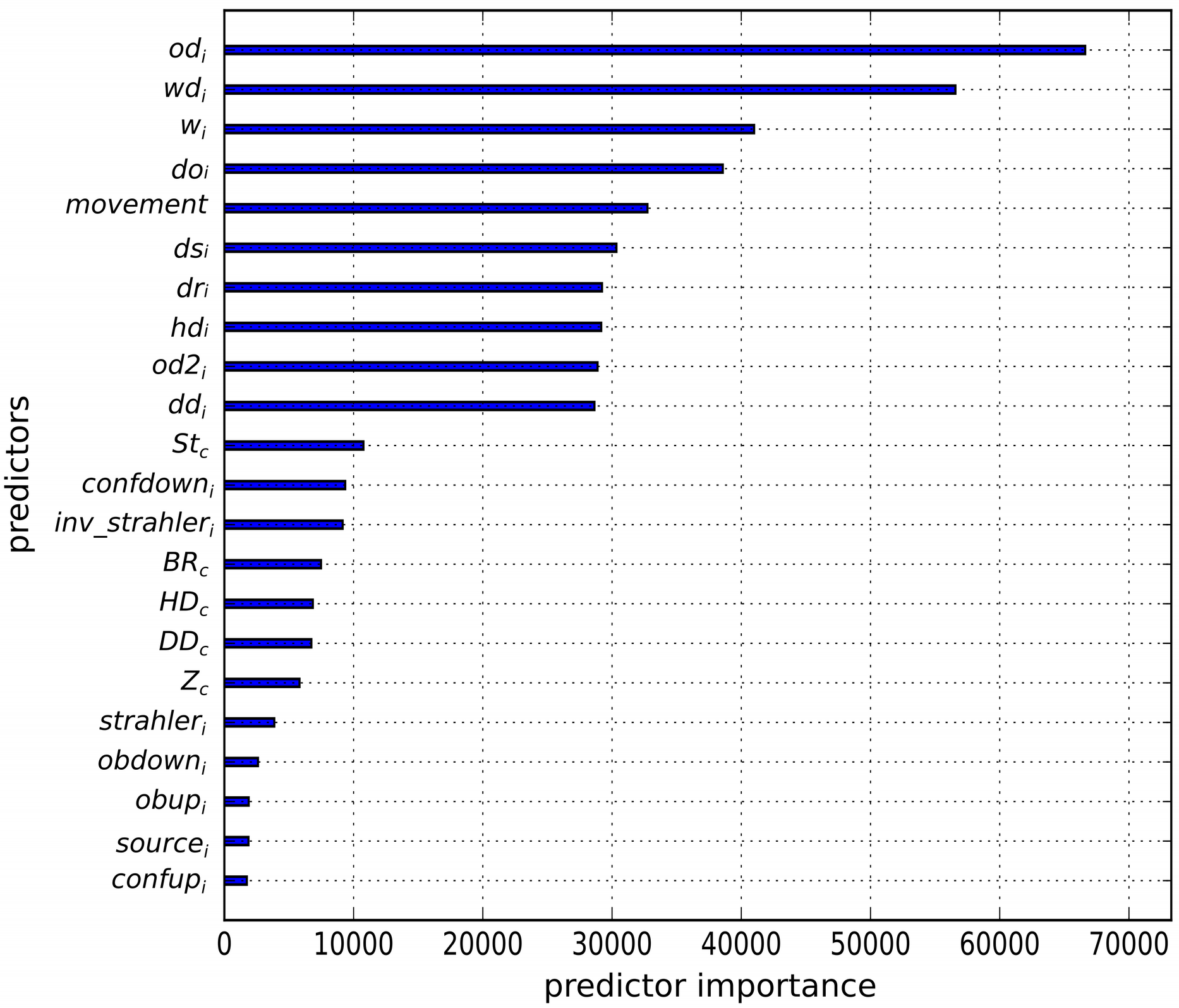
Importance of the different predictors (Table 2) in the BRT for eel density per reach.

Partial dependence plots illustrate marginal effects of one or several predictors of the BRT on the output (Fig. 8). Unsurprisingly, partial dependence plots show that log densities decrease linearly with the distance to the outlet, as expected with a diffusive process. However, it shows, that the densities at the outlet are higher with a flow-wise scenario but decrease faster than with a path-wise scenario. That confirms that a flow-wise scenario produces higher densities close to the sea, and lower densities in upstream parts of river catchments. Partial dependence plots also show that densities decrease with increasing density of obstacles, whatever the scenario.

**Fig. 7.**
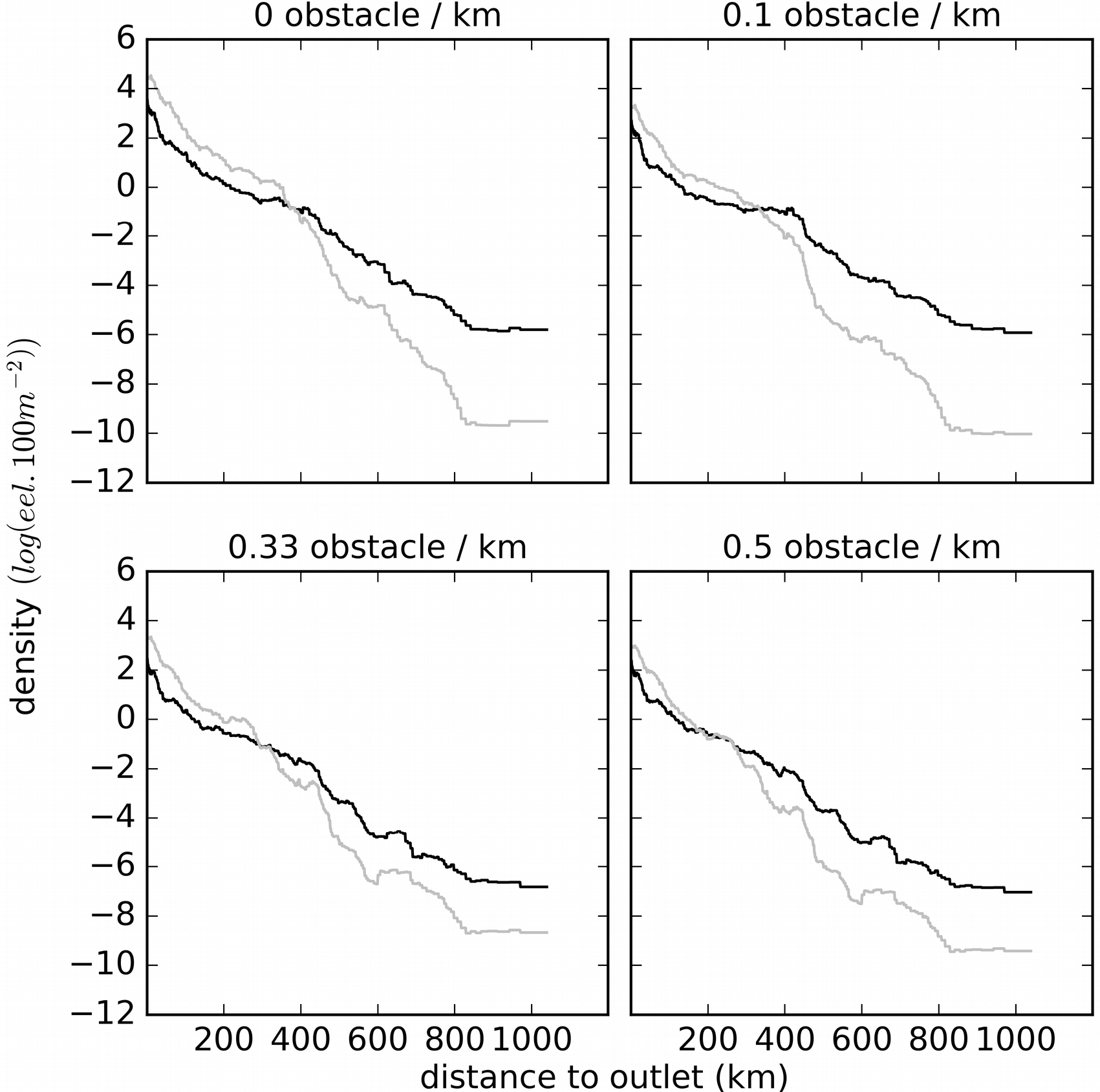
Partial dependencies plots illustrating the effect of *do*_*i*_ on the predicted log density (first BRT). The grey line corresponds to predictions of the BRT with a flow-wise scenario, the black line correspond to a path-wise scenario. Each panel corresponds to predictions with a specific value of *od*_*i*_. Predictions were made with continuous variables set at their average values, EMU “Garonne-Dordogne-Charente-Seudre”, and indicator attributes (*source*_*i*_, *outlet*_*i*_, *obup*_*i*_, *obdown*_*i*_, *confup*_*i*_, *confdown*_*i*_) set to false.

The second BRT that focuses on the discrepancies between the path-wise and the flow-wise scenarios provide similar results (Fig. 9). Hydraulic densities (*hd*_*i*_) and density of drainage (*dd*_*i*_) seem to be slightly more important. This would be logical since these two factors are related to the number of confluences and the behavior at confluence is the main difference between two scenarios. Once again, the discrepancies are mainly influenced by reach attributes, by the number of obstacles to pass and by network configuration while catchments attributes have a more limited effect.

**Fig. 9.**
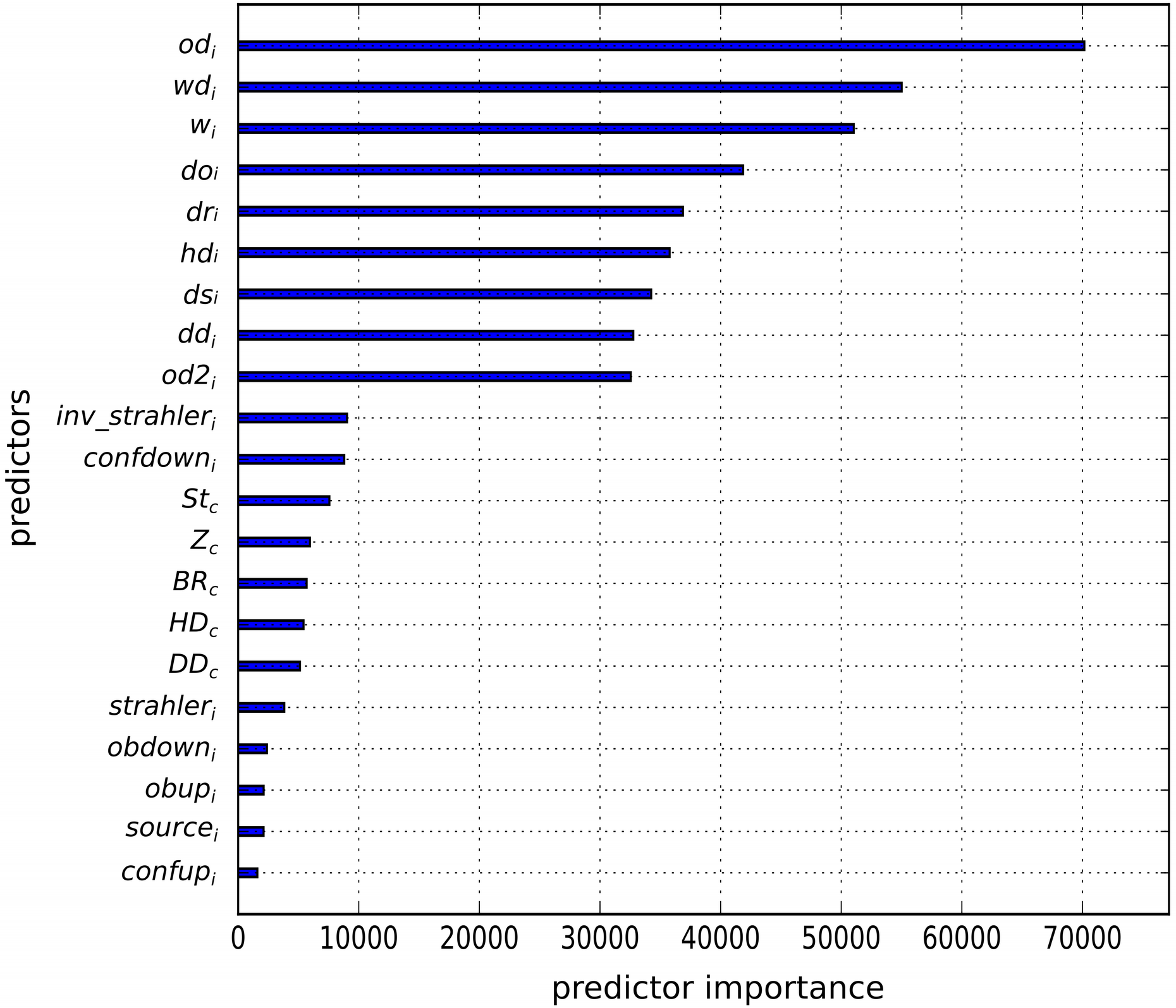
Importance of the different predictors (Table 2) in the BRT for eel density per reach.

## 4 Discussion

### 4.1 Flow-wise or path-wise diffusion: ecological interpretation

This work was conducted with the aim of improving knowledge of the eel colonization process and the effect of human-induced obstacles. Since the study conducted by Ibbotson et al. (2002), eel colonization has generally been assumed to result from a diffusion process. However, diffusion in a dendritic and fragmented river network is not a straightforward process, and flow-wise diffusion and path-wise diffusion represent two plausible mechanisms. Johnson et al. (1995) suggested that path-wise diffusion is more likely for riparian animals, since they are not impacted by water flow. Path-wise diffusion would be typical of random movements dedicated to resource finding, i.e., ranging. Conversely, flow-wise diffusion would correspond to a rheotactic-oriented behavior. Our results demonstrate that path-wise diffusion is more likely than flow-wise diffusion for eels. However, eel colonization is the result of two phases. In the first phase, young eels (glass eels and elvers) display active migrant behavior (Bureau du Colombier et al., 2008; Imbert et al., 2010). This migration is expected to be oriented by different factors including flow direction (Trancart et al., 2014) or intra-specific interactions (Bureau du Colombier et al., 2008; Geffroy and Bardonnet, 2012). In the second phase, eels tend to settle and to display sedentary behavior (Imbert et al., 2010) in which movements are less oriented and would correspond to ranging. While the first phase corresponds to flow-wise diffusion, the second phase corresponds to path-wise diffusion, and our results demonstrate that the resulting distribution of eels in a river catchment after several years in continental waters is driven more by the second phase than the first phase.

### 4.2 Consequences regarding eel distribution and the impact of obstacles

Our results suggest that eel distribution is the combined result of the river network configuration, the position of obstacles and the movement process. This situation leads to very different impacts of obstacles depending on the EMU and scenario, both during the diffusion phase and subsequently during downstream migration. The effect of passing obstacles is not neutral and can have a wide variety of impacts. Direct mortality induced by turbines during downstream migration has been widely studied (Boubée and Williams, 2006; Winter et al., 2007; Calles et al., 2010; Calles et al., 2012). Nevertheless, obstacles can exert a large variety of impacts in both upstream and downstream passages, such as stress, disease, energetic costs (Budy et al., 2002; Garcia De Leaniz, 2008), overpredation (Agostinho et al., 2012) or overfishing (Briand et al., 2003), which can result in additional direct or delayed mortality. Kuby et al. (2005), O’Hanley and Tomberlin (2005) and Palmer and Bernhardt (2006) demonstrated that the dendritic nature of hydrographic networks makes the responses to multiple perturbations more complex and difficult to predict. Van Looy et al. (2014) highlighted the importance of considering both structural and functional connectivity when prioritizing management actions to restore riverine continuity. Our results are consistent with their statement: the impact of obstacles on the fish distribution is the result of fish movement processes, dendritic river network configurations and obstacle locations.

### 4.3 Limits of a random-walk model

The model we developed is based on a simple random walk with a constant transition matrix, implying several strong assumptions. First, in the absence of more precise information, we assumed that all obstacles exhibit similar passability, which is clearly a rough approximation, since the height of obstacles can range from less than a meter to tens of meters. The collection of more precise information on dams, such as the information proposed by Tremblay et al. (2016), would provide a valuable source of information in the future. There is an ongoing initiative with this aim in France, where the goal is to provide the information required to calculate the ICE index (Baudoin et al., 2014) in the ROE database in the future.

Another assumption is that fish movements are only impacted by distance (through elevation to the power of the transition matrix) and obstacles (through the passability coefficient), although other abiotic factors are likely to play a role. For example, no eels are observed at an altitude superior to 1000 m in France (Briand et al., 2008). Salinity, temperature, habitat quality and mortality are also likely to influence the probability of transition. However, many of these factors are highly correlated, with altitude being lower in downstream parts of river catchments, while salinity increases in estuarine areas. Moreover, the data available for downstream parts of catchments are scarcer because electrofishing sampling is difficult to perform in deep zones. In the future, it would be interesting to incorporate additional sources of data, such as sampling results from the Water Framework Directive monitoring program or the Eel Management Plan monitoring programs. Despite being based on different sampling protocols and, thus, requiring intercalibration, the results of such monitoring programs conducted in deeper habitats should help to disentangle the effects of other factors related to accessibility. The amount of available habitats is also likely to influence fish movements and settlement. In the future, it would be interesting to incorporate local habitat quality in the model by coupling TABASCO with habitat models that predict habitat suitability for fishes (Lamouroux et al., 1998; Lamouroux and Jowett, 2005).

A main drawback of random-walk is that it leads to many round-trips around obstacles. In real world, fishes probably avoid round-trips and it would be interesting to take into account the possible evolution of passability through iterations because of some despondency. Moreover, we did not take into account any mortality during passage. However, mortality occurs during upstream passage because of predation (Agostinho et al., 2012) or fishing (Briand et al., 2003). Mortality also occurs during downstream passage, for example because of passages through hydropower plants (Boubée and Williams, 2006; Winter et al., 2007; Calles et al., 2010; Calles et al., 2012). Even if such mortality rates are low, a significant number of eels will be killed given the number of passages.

This mortality should be accounted for in the future. Additionnally, we use a random-walk to mimic a diffusion process, however, such a diffusion process at the population scale can arise from different processes at the individual scale (Okubo and Levin, 2013). In a random walk, every fishes moves at each time step. But a diffusion process can also arise from individual movements down densities gradients: at each time step, movements occur only from habitats with higher densities towards habitats with lower densities and only a fraction of individuals move. Assuming such a movement would have led to the same overall result at the population scale but would have reduced the number of obstacles passages.

### 4.4 Potential improvements

Another strong limit is that the model is currently fitted on all yellow eels with total lengths of 150 mm to 900 mm, though these eels do not necessarily have the same behavior (Imbert et al., 2010). This is clearly a strong assumption, since the population and recruitment are declining (Dekker, 2000; Drouineau et al., 2016). Moreover, growth rates are highly variable within catchments (Daverat et al., 2012; Helfman et al., 1984; Melià et al., 2006), with the growth phase lasting much longer in upstream freshwater habitats than in downstream brackish waters. These ecological issues and spatial heterogeneity in anthropogenic pressures affect yellow eel indices, and the Working Group on Eels in charge of yearly assessments of European eels (the ICES (International Council for the Exploration of the Sea) -EIFAAC (European Inland Fisheries and Aquaculture Advisory Commission) -General Fisheries Commission for the Mediterranean (GFCM) working group) identified more significant trends in recruitment or escapement indices than in yellow eel indices, which are often very flat (ICES, 2012). Age data would enable the model to be fitted to cohorts, but such data would require the development of an otolith reading program at a very large scale. As a last resort, the model may also be fitted to length classes, although growth rates are highly variable between habitats types, which may limit the variability of eels. Such distinctions between age or length classes would be even more important to account for habitats quality within the model, as discussed earlier, since eels of different sizes display different behaviours.

Another consequence is that we mixed young eels exhibiting migrant behavior with older eels exhibiting diffusive behavior (Imbert et al., 2010). Young eels are smaller and consequently exhibit lower catchability and are therefore probably underestimated by electrofishing. Moreover, in the context of declining populations (Drouineau et al., 2016), recruitment is becoming increasingly lower, and the proportion of migrant individuals in a population decreases over time, which may partly explain why the path-wise model provided a better fit to the data. It would be interesting to fit the model to the smallest eels to determine whether the flow-wise scenario, which is more likely in a context of upstream migration, provides a better fit to the data than a path-wise scenario, which is more likely in a dispersive population.

### 4.5 The relevance of a mechanistic model of colonisation fitted by maximum likelihood

The model was fitted by maximizing the log-likelihood function, which is a classical approach (Bolker et al., 2013) with several advantages: it provides a unique solution and converges asymptotically to an unbiased solution. Moreover, by inverting the Hessian matrix at the optimum, we obtained the variance-covariance matrix of estimated parameters and were able to build confidence intervals. However, in the context of the present study, one of the parameters was an integer (*s*). It raises problems to compute the derivatives of the likelihood function with respect to this parameter and consequently the Hessian matrix and the confidence intervals should be taken with caution. That probably explains some of the strong correlations between parameters estimates. Additional work will be required to overcome this limitation, but this question of confidence intervals for a discrete parameter is not a straightforward problem (Choirat and Seri, 2012). Moreover, in the optimization process, we considered s as a continuous parameter. In the future, it would be interesting to use a mixed-integer non-linear optimization algorithm to achieve more appropriate optimization.

To date, two main approaches have been used to address the question of the impact of fragmentation on the distribution of eel or diadromous species at large scales. Purely statistical models have been developed for eels. For example, Eel Density Analysis (Briand et al., 2015) is based on a generalized additive model fitted to electrofishing data, using various ecological and anthropogenic factors, including human-induced obstacles as explanatory variables. For lampreys and shad, Segurado et al. (2014) developed species distribution models to assess the impact of obstacles on the distribution of fishes. Statistical models have many advantages, including their flexibility and the possibility of using them even in the absence of precise knowledge of the underlying mechanism. Another approach in landscape ecology is based on the use of structural connectivity indicators according to graph theory (Pascual-Hortal and Saura, 2006; Segurado et al., 2013). However, these indicators are generally based on assumptions regarding fish movements among patches, and methods for checking those assumptions are lacking. In this context, TABASCO fills a gap in the range of available tools, providing a mechanistic model that is complementary to statistical models and suitable for verifying the relevance of graph-based indicators. In the context of climate change, the impact of global warming on the distribution of fishes is mainly being evaluated with statistical species distribution models, and there is a call for mechanistic models (Franklin, 2010; Keith et al., 2008; Thuiller et al., 2008) to achieve better predictions by accounting for the species dispersion capacity. The development of such mechanistic models has been demonstrated to be very useful in providing congruent results to correlative approaches (Kearney et al., 2010; Rougier et al., 2015). We are convinced that the same statement is valid for modeling the impact of fragmentation in dendritic river networks and that such mechanistic models can be used to support decision making in the prioritization of connectivity restoration actions (Drouineau et al., 2018).

In this study, we applied TABASCO to the European eel in France. However, the model can be easily applied in other countries and to other temperate eels because the assumptions and data requirements are limited. Theoretically, the model can also be applied to any freshwater species displaying diffusive behavior in continental water, although the indicator vector *Ic* should be modified to account for multiple sources of dispersion.

## 5 Acknowledgements

This study was funded by the Office National de l’Eau et des Milieux Aquatiques (Onema) / Agence Française de la Biodiversité (AFB). We are very grateful to Cédric Briand for his tremendous help during this project. We would also like to thank Herve Pella for his support with the RHT layer.

